# Expression of extracellular Hsp90 is a molecular signature of T cell activation, providing a means to image and target T Cell activation in autoimmune disease

**DOI:** 10.1101/2021.06.06.446823

**Authors:** Scott A. Scarneo, Aaron P. Smith, Jacob Favret, Robert O’Connell, Joy Pickeral, Kelly W. Yang, Guido Ferrari, David R. Loiselle, Philip F. Hughes, Manjusha M Kulkarni, Madhusudhana Gargesha, Bryan Scott, Debashish Roy, Barton F. Haynes, Jesse J. Kwiek, Timothy A. J. Haystead

**Affiliations:** Department of Pharmacology and Cancer Biology, Duke University School of Medicine, Durham, NC 27710; Bolder BioPath INC., Boulder, CO, 80301; Department of Surgery, Duke University School of Medicine, Durham, NC 27710; Department of Microbiology, Ohio State University, Columbus, OH 43210; BioInVision, Inc., Mayfield Village, OH 44143; Department of Medicine, Duke University School of Medicine, Durham, NC 27710

**Keywords:** T cells, Heat shock protein 90 (Hsp90), tethered inhibitor of Hsp90, eHsp90, CD3 and CD28 antibodies, CD69, CD25, rheumatoid arthritis, inflammatory bowel disease, autoimmunity

## Abstract

Heat shock protein 90 (Hsp90) maintains cellular proteostasis during stress and has been under investigation as a therapeutic target in cancer for over two decades. We and others have identified an extracellularly expressed form of Hsp90, eHsp90, that previously appeared to be restricted to rapidly proliferating cells exhibiting a metastatic phenotype. Here, we used HS-131, a fluor-tethered eHsp90 inhibitor, to quantify the effect of T cell activation on the expression of eHsp90 in human and mouse T cells. In cell based assays, stimulation of human T cells induced a 20-fold increase in eHsp90 expression at the plasma membrane, suggesting trafficking of eHsp90 is acutely regulated by TCR and inflammatory mediated signaling. Following injection of HS-131 in mouse models of human rheumatoid arthritis and inflammatory bowel disease, we detected localization of the probe at sites of active disease, consistent with immune cell invasion. Moreover, despite rapid hepatobiliary clearance, HS-131 demonstrated efficacy in delaying the onset and progression of disease in the arthritis model. Our results suggest eHsp90 expression at the plasma membrane of T cells is a molecular marker of autoimmune induced activation and potentially a therapeutic target for chronic diseases such as rheumatoid arthritis and inflammatory bowel disease.

**Perspective:** T cells mediate many maladaptive disease pathologies such as autoimmune disorders. Many immunosuppressants in clinical use act by preventing the activation of T cells. However, this approach can lead to increased rates of opportunistic infection and malignancy. Here, we show for the first time that eHsp90 is upregulated during T cell activation in response to exogenous ligands. Therefore, pharmacological inhibition of eHsp90 may provide a means to target only activated T cells for immunosuppression. Selective targeting of only activated T cells could be efficacious against autoimmune disease without increasing the incidence of undesirable outcomes. Additionally, the presence of eHsp90 on the surface of activated T cells could be diagnostically useful to track the status and progression of autoimmune disease in patients.

## Introduction

T cells play an integral role in the host adaptive immune response, choreographing immune surveillance and the host response to infection [13]. Following T cell receptor (TCR) stimulation, T cells undergo rapid replication and activation, and respond to their target antigens by secreting immunoregulatory cytokines and chemokines or through direct cell killing [11]. These rapid cellular changes are governed by TCR mediated protein kinase signal transduction, which increases the expression of specific CD markers essential to the T cell response [10; 21]. Heat shock protein 90 (Hsp90) and some of its co-chaperones are upregulated during T cell activation and have been shown to play an important role in regulating many protein kinases essential for T cell activation [8; 18; 19]. Hsp90 inhibition has also been shown to block T cell function i*n vivo*, as seen in mouse models of allograft rejection and in rheumatoid arthritis [16]. Thus, Hsp90’s role in mediating T cell proliferation is essential to T-cell mediated immune surveillance.

Recently, we and others characterized an extracellular form of Hsp90 (eHsp90) that appeared to be only expressed on the surface of rapidly proliferating tumor cells exhibiting a metastatic phenotype [3; 7; 23],[6; 25]. This phenomenon was originally characterized by Neckers and colleagues, showing that antibodies to Hsp90 block cell migration in tumor lines expressing the surface form of the protein [24]. Subsequently, using cell impermeable fluor-tethered inhibitors of Hsp90, we utilized confocal microscopy to study the trafficking of eHsp90 on the surface of various aggressive tumor cell lines [3],[7]. These studies revealed that the protein was actively trafficked to the plasma membrane (PM) under oncogene control and upon binding of a tethered Hsp90 inhibitor, formed aggregates (90 bodies) that were actively reinternalized [7]. *In vivo* studies with tethered Hsp90 inhibitors in mice bearing breast tumors showed expression of eHsp90 was indeed exclusively associated with malignant tumor cells [7]. Subsequently, a tethered inhibitor of Hsp90 carrying a far red fluorophore (HS-196) has entered phase 1 clinical trials as a diagnostic of tumor malignancy.

Here, we show that activated CD^69+^/CD^25+^ T cells express high levels of eHsp90 at the plasma membrane (PM) in response to CD3/CD28 antibody challenge, suggesting that trafficking of eHsp90 is a regulated process in T cells and not a phenomenon of cellular stress such as occurs in oncogenic transformation. Expression of the protein in T cell populations was followed both *in vitro* and *in vivo* with HS-131, a Cy5 carrying tethered inhibitor of Hsp90. Furthermore, systemic evaluation of eHsp90 expression in murine models of autoimmune disease shows that tissues with robust expression of eHsp90 also show histopathological evidence of active disease. Our results show that targeting of eHsp90 provides a selective diagnostic means to evaluate active disease sites in autoimmunity and suggest that tethered inhibitors of Hsp90 could be developed as a means to achieve precision immunosuppression *in vivo*.

## Results

### Regulation of Hsp90 chaperone machinery in T cells

Using a list of 140 known genes that comprise the entire cellular chaperone machinery we interrogated publicly available genome wide RNA-seq data derived from activated human CD4^+^/CCR6^+^ human peripheral blood mononuclear cells (PBMC)’s ^12^. Expression of both isoforms (α and β) of Hsp90 were highly upregulated over the time-course of activation, along with other stress induced chaperones including Hsp70, Hsp60, and Hsp40 (Fig.1A, Supplemental F1). We also found large expression changes in several Hsp90 co-chaperones including AHSA1, FNIP2, STIP1, FKBP4, PPID, and PP5 ^13^. Others such as FNIP1, STUB1, and FKBP5 were reduced in expression. We confirmed the increased expression of known activating and regulatory proteins of Hsp90 such as FKBP4/5, Trap1, Hop, Chip, and Aha1 by western blot. Consistent with the RNA-seq data, these regulatory and co-chaperone proteins were significantly upregulated at 72 and 96 hours post stimulation compared to resting T cells. The addition of exogenous IL-2 at 72 hours post stimulation did not affect these changes in expression. (Figure 1B). Consistent with expression analysis, activated human PBMC T cell populations showed a marked induction of both Hsp90α and Hsp90β protein by 3 and 4.5-fold respectively (p<0.001) (Figure 1c). These findings are consistent with a growing body of evidence that Hsp90 is necessary for T cells to respond to immune or cytokine challenge ^2,4,5,7^.

**Figure 1.**
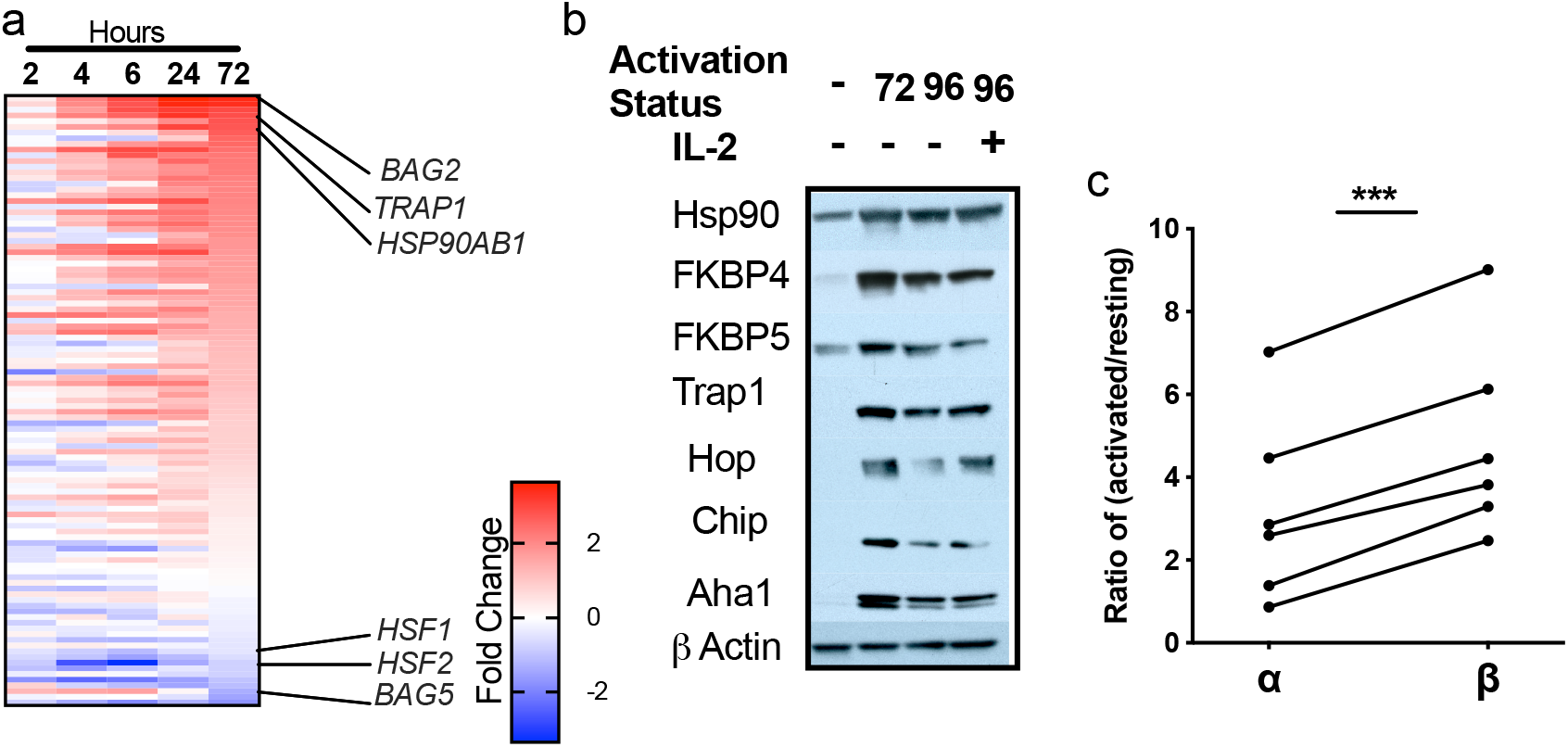
Hsp90 upregulation following T cell activation. (A) mRNA expression of co-chaperones and mitochondrial homologs of Hsp90 72 hours following T cell activation. (B) T cell isolated from PBMC’s were activated with anti-CD28 and anti-CD3 antibodies for 0, 72 or 96 hours with or without additional IL-2 stimulation and Hsp90, co-chaperones and mitochondrial homologs were probed via western blot for relative protein expression. (C) Hsp90 isolated from T cells 24 hours post activation and purified on a Hsp90 specific resin analyzed by MALDI mass spectrometry, for alpha and beta Hsp90 isoforms. Data represents 6 biological replicates. The data was analyzed with paired t-test. ***p<0.001.

A hallmark of T cell function is to proliferate rapidly in response to antigen presentation along with appropriate cytokine and costimulatory signaling. This leads to the induction of the chaperone machinery which is essential in facilitating this process. Data shown in Figure 1C shows a marked induction of Hsp90α/β at the protein level. In some respects, antigen induced T cell proliferation may be analogous to oncogene driven metastasis. Therefore, to investigate whether proliferating T cells similarly express eHsp90 during T cell activation we examined their ability to absorb fluor-tethered Hsp90 inhibitors following an anti-CD3/CD28 challenge (Fig 2). Figure 2a shows the structures of two types of fluor-tethered Hsp90 inhibitors used in our studies. The probes consist of an Hsp90-binding ligand, a polyethylene-based tether (linker) and a fluorophore (Cy5 dye ex640nm/em680nm). Both the ligand and the fluorophore moieties can be substituted with a variety of small molecules. For example, in our studies we synthesized 3 types of Cy5 probe in which the ligand was either substituted to an N’N dimethylamide to create an inactive control molecule (HS-198) or equally potent but cold-soluble (4°C) analog by substituting the benzylamine (HS-131) with a benzamide moiety (HS-132). Flow analysis of human PBMCs treated with or without CD3/CD28 antibodies showed dose dependent uptake of HS-132 in the activated T cell population, in contrast to resting T cells (Fig. 2b). When the study was repeated at 4°C versus 37°C, uptake of the probe was blocked at 4°C, suggesting that uptake involves active cellular processes rather than simple diffusion (Fig.2c). Similar temperature dependent effects were observed previously with malignant tumor cell lines [7]. Figure 2d shows the uptake of the probe is eHsp90 specific, since minimal uptake was observed with the HS-198 inactive analogue. Additionally, the fluor signal was effectively blocked by competition with a saturating dose of a structurally non-related ATP competitive inhibitor of Hsp90, ganetespib [12; 26].Therefore activated T cells specifically internalize fluor-tethered Hsp90 inhibitors due to interactions with the ATP binding site of eHsp90 expressed at the plasma membrane. Confocal analysis of isolated activated human T cells confirms uptake of HS-132 following CD3/28 stimulation compared to resting (inactivated) T cell populations (Fig.2e). Furthermore, the binding of HS-132 was eliminated by HS-10, a non-fluorescent Hsp90 inhibitor. (Fig.2e).

**Figure 2.**
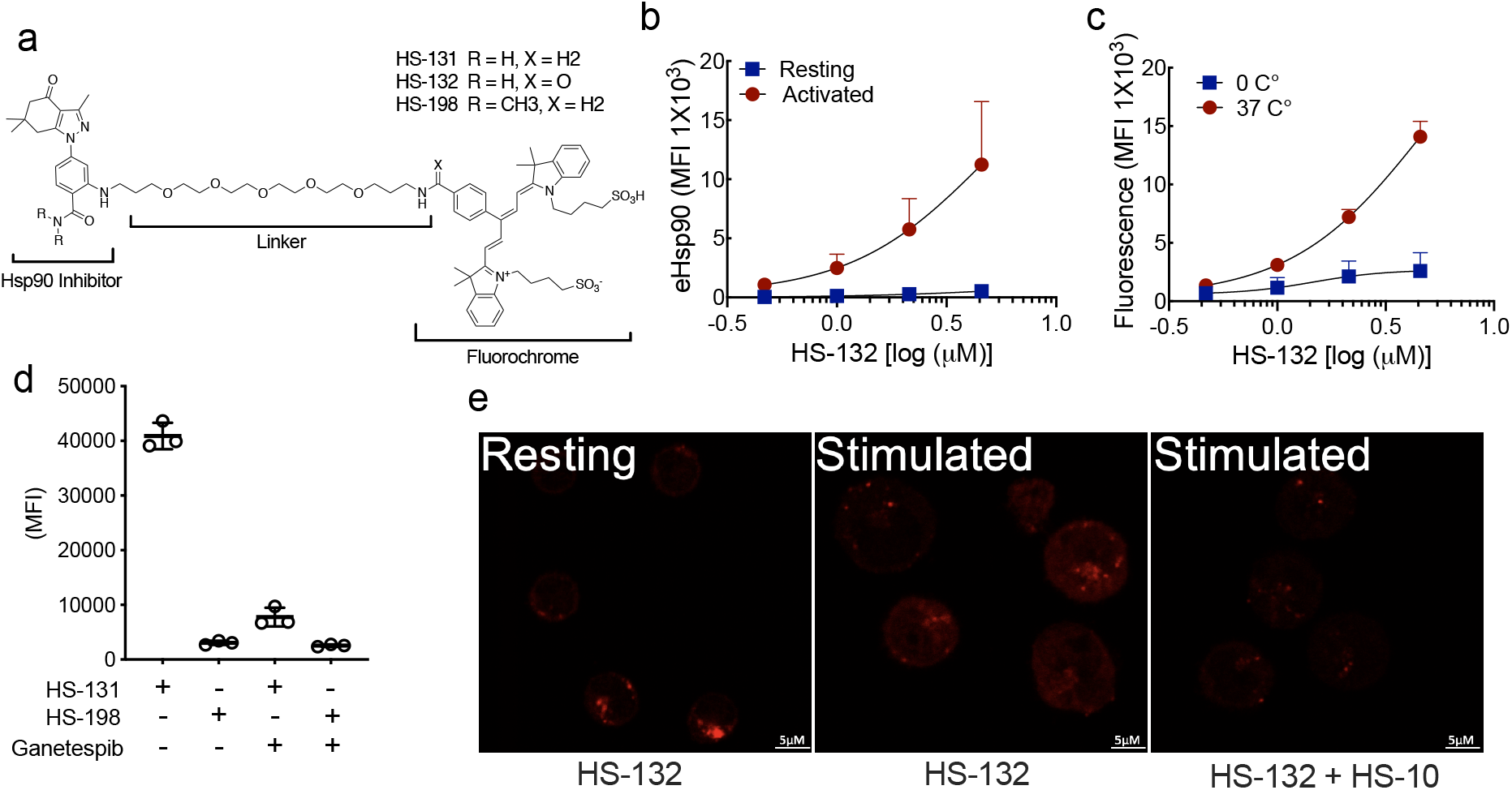
T cell activation induces extracellular Hsp90 (eHsp90) expression. (A) Chemical structure of HS-131, and HS-132 active molecule, the inactive analogue HS-198 (negative control). (B) Naïve T cells isolated from healthy human peripheral blood mononuclear cells (PBMC’s) were activated with anti-CD28 and anti-CD3 antibodies, or resting (unstimulated) for 96 hours and treated with HS-132 at the indicated concentrations. Data represented as mean± SD, 6 biological replicates. (C) Internalization of probe by T cells at physiological (37°C) and non-physiological (4°C) temperatures at varying concentrations of HS-132. Data represents mean± SD, 4 biological replicates. (D) T cells isolated from stimulated human PBMC’s were treated with 10µM HS-131 (active molecule) and HS-198 (inactive control) ± a 10 fold excess of ganetespib. Data represents mean± SD, 3 biological replicates. (E) Confocal analysis of HS-132 uptake in stimulated T cells, which is blocked with treatment of the non-fluorochrome tethered Hsp90 inhibitor HS-10.

We next sought to track the expression of eHsp90 and total Hsp90 over the course of T cell activation. We evaluated both total Hsp90 cellular content and eHsp90 expression in anti-CD3/28 stimulated human PBMCs. Total Hsp90 expression is upregulated 5-7 fold in CD3+ T cell at 2 to 3 days post activation, followed by a reduction in Hsp90 expression. In contrast, eHsp90 expression, as determined by HS-131 binding, was upregulated 20-25-fold following activation, which is significantly more than the observed change in total Hsp90 expression (Figure 3a). Specifically, anti-CD3/28 stimulation caused eHsp90 up regulation in CD69+/CD25+ and CD69+/CD25-T-cell populations with a 5-8 fold increase within the first 24 hours post activation, whereas CD69-/CD25-T cells show no increase in probe or eHsp90 levels (Figure 3b). Total Hsp90 was also upregulated in the CD69-/CD25+ populations but only by approximately ∼2.5 fold (Figure 3c). When we blocked eHsp90 entry with monoclonal Hsp90 antibodies, no change in viability, the percentage of CD25+ cells or even overall cell numbers was observed, indicating blocking of eHsp90 internalization does not interfere with T-cell activation (Figure 4s). This data suggests that eHsp90 may function in the delivery of client proteins to the plasma membrane. One candidate client eHsp90 protein is CD25, the IL-2 receptor. This protein is highly upregulated during T-cell activation due to a positive feedback loop mediated by autocrine and exocrine IL-2^18^. Here we observe that CD25 expression peaks at 72 hours post activation, followed by a reduction in expression thereafter which correlates with reduced eHsp90 expression (Figure 3d). Due to the tight correlation of CD25 and eHsp90 expression, we further tested whether exogenous IL-2 stimulation could maintain high expression of both proteins in activated T cells 96 hours post-stimulation. We found that CD25 expression is significantly upregulated along with eHsp90 in a dose dependent manner in response to IL-2 (Figure 3e, Supplemental 2s). Total cellular Hsp90 expression was not affected by IL-2, suggesting that the trafficking of eHsp90 to the cell surface in T cells is part of a receptor mediated process rather than a reflection of changes in overall Hsp90 protein expression.

**Figure 3.**
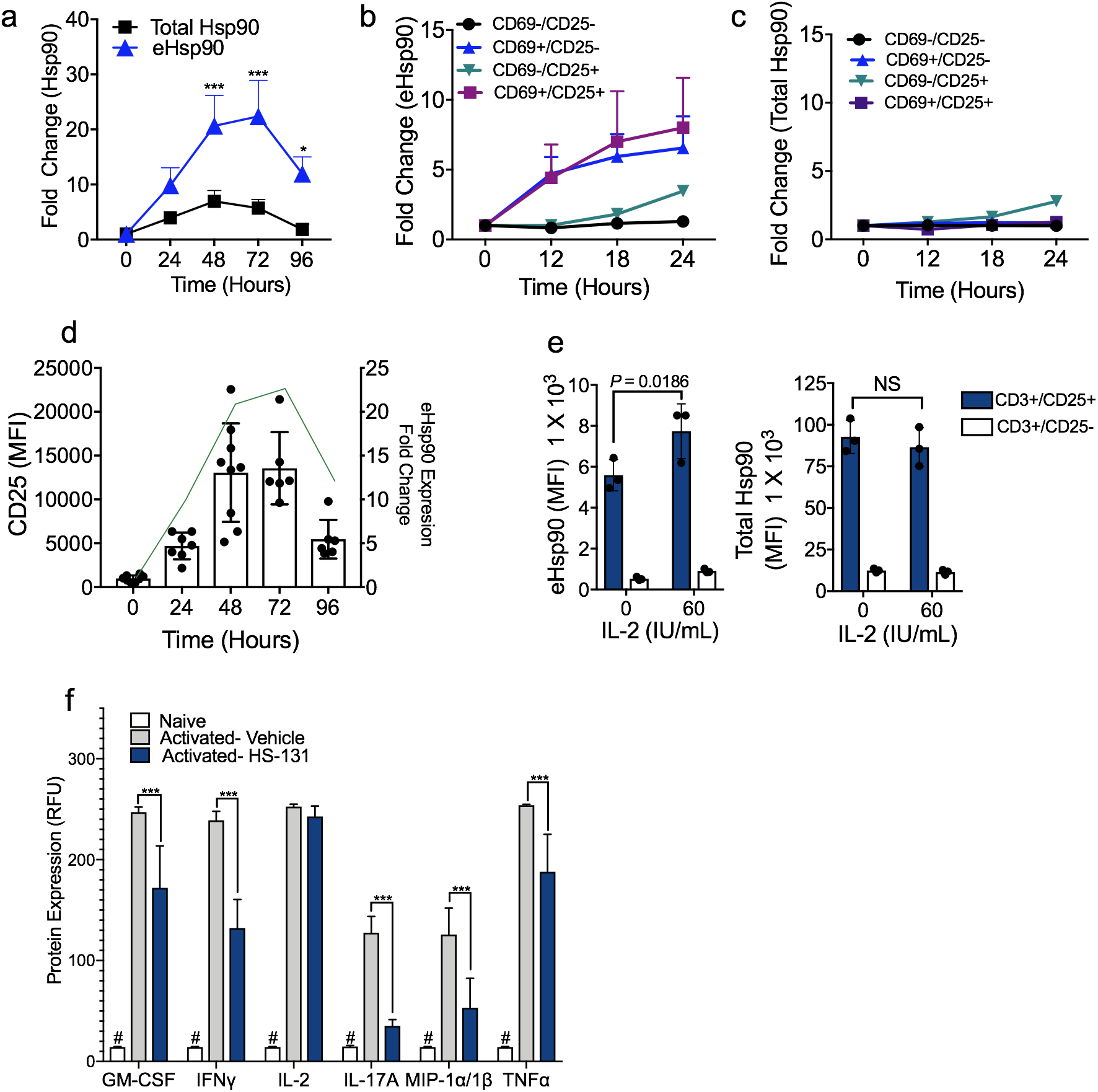
(A) Temporal upregulation of eHsp90 compared to total Hsp90 levels in T cells stimulated with CD3/CD28 antibodies over a 96-hour period. The data were analyzed by 2-way ANOVA with Sidak’s multiple comparison test, * p <0.05, **p<0.01, ***p<0.001. Experiments were performed with 6 biological replicates. (B) Flow cytometric analysis of CD3, CD25, and CD69 T-cell populations relative eHsp90 expression compared to baseline. (C) Upregulation of total Hsp90 in activated T cell populations. Experiments were performed with 3 biological replicates for each timepoint (b,c). (D) Mean (±SD) CD25 expression on T cells 24-96 hours post CD3/28 stimulation. Temporal upregulation of eHsp90 (green line) post T cell stimulation. CD25 expression; 6-8 biological replicates per time point. eHsp90 expression 6 biological replicates for each timepoint. (E) eHsp90 and total Hsp90 expression in CD3+ T cells isolated from PBMC’s were stimulated with IL-2 (60IU/mL) for 24 hours. Data represent mean±SD. The data were analyzed by 2way ANOVA with Sidaks multiple comparison test. Experiments were performed once with 3 biological replicates. (F) Cytokine expression 24 hours post CD3/28 T cell stimulation treated with or without HS-131 (1μM). Data represent mean±SEM. The data were analyzed by 2way ANOVA with Sidaks multiple comparison test. Experiment was performed once with 4 biological replicates.

**Figure 4.**
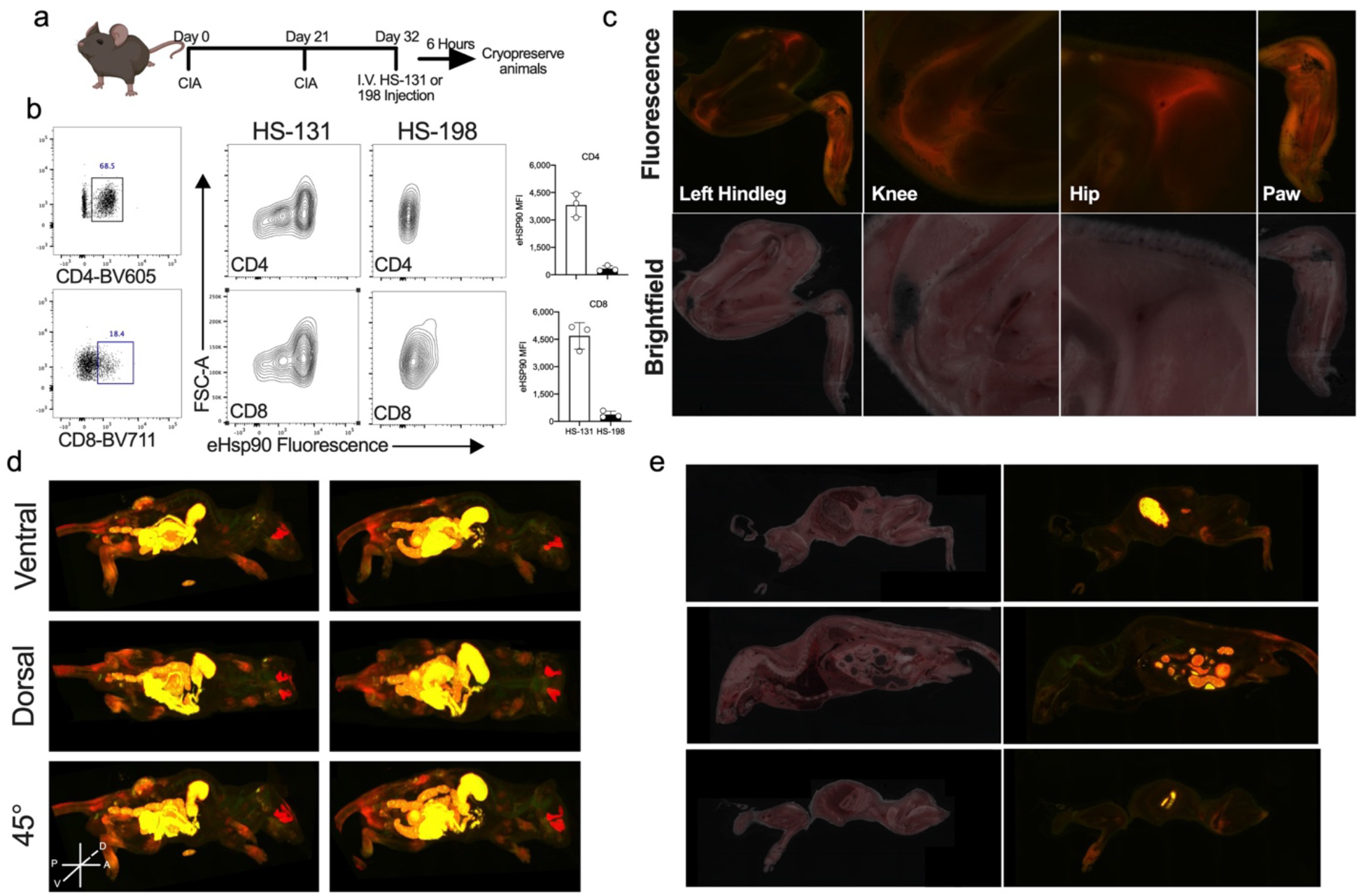
(A) Schematic of CIA mouse model of RA experimental design, mice were inoculated with CIA on day 0 and 21. At peak of disease onset (day 32) animals received I.V. Hs-131 or HS-198 (inactive control). 6 -hours post treatment mice were whole body cryopreserved. (B) Flow cytometric analysis of eHsp90 expression on CD4 and CD8 T Cells isolated from the serum of HS-131 and HS-198 treated mice 6 hours post tail vein injection. Gated regions indicate CD4 and CD8 T -Cell populations isolated from individual mice. eHsp90 expression in CD4 and CD8 T cells 6 hours post HS-131 or HS-198 treatment. N=3±SD. Experiments represent 3 biological replicates (C) Brightfield and fluorescent histology of hindleg, knee, hip and paw of CIA mice 6 hours post tail vein injection of HS-131. (D)Whole body reconstruction of individual HS-131 treated CIA mice 6 hours post tail vein injection. (E) Brightfield and fluorescent cross section of CIA mouse.

### *In vivo* tracking of eHsp90 expression in autoimmune disease states

Previous work suggested that the expression of eHsp90 was restricted to tumor cells exhibiting a metastatic phenotype. This was consistently seen in cell culture, tumor cell xenografts and mouse models of spontaneous metastatic disease [3]. Our studies with isolated human PBMCs now suggest that activated T cells also upregulate eHsp90 in response to pro-inflammatory activation. To determine if this phenomenon occurred within the context of a fully functional immune system we utilized the collagen induced arthritic (CIA) mouse model of human rheumatoid arthritis (RA) [16]. Mice (DBA/1lacJ ∼8 weeks old) reliably develop polyarthritis when immunized against bovine type II collagen[4; 17]. The disease that occurs is usually non-symmetric, and any combination of paws/joints may be affected. Mice were injected via the tail vein with bovine collagen at day 0 and then again 21 days later. This second injection triggered the development of the CIA related symptoms over the next several days within the joints (Fig.4A). A primary mediator of local inflammatory responses in autoimmunity are invading T cell populations recognizing foreign antigens. To determine if invading T cells at sites of active disease expressed eHsp90, we injected the animals with HS-131 or HS-198. This was done at the peak of RA symptoms following the second immune challenge. After 6 hours, the animals were euthanized, immediately cryo-frozen in liquid nitrogen and cryo-sectioned (Fig.4). Approximately, 500 40µm slices were made and each slice imaged with both bright field and by fluorescence at Ex640nm/Em680nm. Evaluation of the biodistribution was then made by histological examination of individual slices and after *in silico* reconstruction of the entire mouse anatomy (Fig. 4 and Supplemental movie 1).

Immune cell phenotyping by flow cytometry of PBMC populations from the CIA mice showed pronounced eHsp90 expression across T-cells as measured by uptake of HS-131. For example, lymphocyte populations of inflammatory CD4 and CD8 cells both showed null, medium and high eHsp90 expression indicative of varying activation states *in vivo* (Figure 4b, Supplemental 3s). No signal was obtained after flow studies on PBMCs isolated from CIA mice treated with HS-198, further showing the specificity of HS-131 for eHsp90 *in vivo* (Fig 4b insert). Histological examination of joint slices taken from the cryo-sectioned CIA HS-131 mouse showed enhanced fluorescence in the joints of the animals, which also correlated with mean arthritic edema scoring and anatomic changes associated with local edema. In particular, the hind leg, knee, hip and paw all showed a marked increase in HS-131 localization, congruent with infiltration of significant lymphocyte and leukocyte populations (Figure 4c). Analysis of the whole-body reconstruction of CIA mice shows systemic HS-131 distribution is largely confined to areas of active immune cell invasion such as the hind paws and joints of the animals (Supplemental movie 1). No HS-131 uptake was found in rapidly dividing progenitor cells derived from epithelial stem cells at the intestinal crypts. The movie and sections shown in figure 4 also highlight the major route of HS-131 elimination which is via the hepatobiliary system and intestines, similar to our prior work in mice bearing breast tumors. Additionally, the fluorescence associated with the eye shown in Figure 4 was also noted in our previous study. This is a natural fluorescence at Ex640nm/Em680nm associated with the Harderian glands found in rodents [22] Importantly, no fluorescence was observed in the joints or paws of the tumor bearing animals in our prior studies [7].

To further illustrate the specificity of HS-131 for sites of active immune cell localization we repeated our cryo-sectioning studies in a mouse model of inflammatory bowel disease (IBD). In this model naïve T cells are isolated and transferred to SCID mice via intraperitoneal injection. Following T cell adoption, the mice develop symptoms of IBD similar to human disease including epithelial hyperplasia, extensive immune cell infiltration and distended colon [15]. Since HS-131 is exclusively eliminated through the biliary system and intestine we waited 24 hours post HS-131 tail vein injection to image mice in an effort to reduce the non-specific intestinal fluorescence signal associated with hepatobiliary probe clearance. Brightfield imaging of IBD mice showed extensive intestinal distention and inflammation consistent with the development of disease. Similarly, extensive fluorescent signaling from the inflamed bowels indicated areas of high eHsp90 expression consistent with infiltrating immune cell populations (Figure 5A and Supplemental movie 2). In comparison, CIA mice 24 hour post HS-131 injection showed no discrete fluorescence associated with the lower intestine (Figure 5b). Comparison of the biodistribution of HS-131 between the two auto-immune models also illustrated the specificity of the probe for sites of immune induced inflammation. In the case of the IBD mouse, no uptake of HS-131 was observed in any of the animal’s joints or paws (Fig.5A and B).

**Figure 5.**
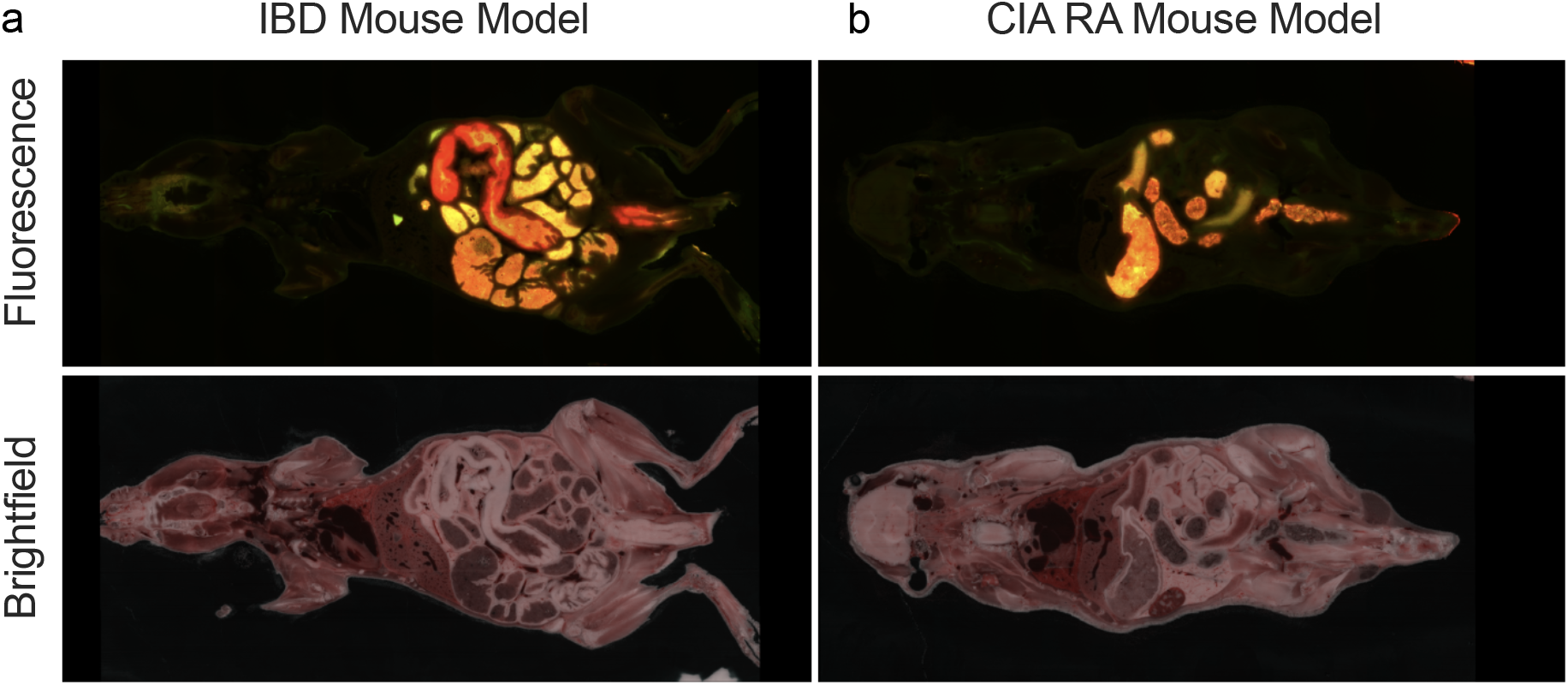
(A) eHsp90 expression in an inflammatory bowel disease (IBD) model of auto immune disease. We utilized the T cell transfer model of IBD to evaluate the imaging potential of HS-131 for other autoimmune diseases. Naïve T cells isolated from female donor mice are transferred into SCID mice via IP injection. 3-5 weeks post transfer mice develop inflammation, an extended colon and epithelial hyperplasia. 24 hours post HS-131 tail vein injection mice were cryopreserved and imaged as previously mentioned. (B) Comparison to CIA mouse 24 hours post HS-131 injection. Experiments represent one mouse.

Previous studies have shown that non-tethered Hsp90 inhibitors may act as anti-inflammatory agents by suppressing pro-inflammatory cytokine production in stimulated immune cells [16]. When administered to animals either orally or by injection, these small diffusible molecules are absorbed in all cell types systemically. In contrast, as shown herein and in prior work with tumor disease models, fluor-tethered inhibitors preferentially accumulate in cells expressing eHsp90. Therefore, to test the therapeutic potential of selectively targeting only eHsp90 as an immune-suppressant we tested HS-131 for therapeutic efficacy in the CIA mouse model. First, we established the *in vivo* bioavailability and pharmacokinetics of HS-131 following i.p. administration. Upon daily dosing, following administration of the second collagen challenge we observed that the pathogenesis of disease development in the HS-131 treated group was delayed compared to the vehicle control (Fig.6a). Specifically, by day 28, 100% of mice in the vehicle cohort had developed significant edema in at least one joint. In contrast, only 75% of the HS-131 treated mice showed disease at day 28 and 100% progression to disease did not occur until day 33. Overall, treatment with HS-131 significantly reduced the mean clinical arthritic score of mice throughout the study period when compared to vehicle control (p<.0053) (Figure 5b). Analysis of the area under the curve showed a ∼23% decrease in clinical score between the groups (Figure 5c). Given the rapid clearance of fluor-tethered versions of Hsp90 inhibitors via the hepato-biliary system, it is remarkable that we observed any efficacy in the CIA model. Our findings suggest that development of longer acting tethered inhibitors of Hsp90 is warranted to achieve a better therapeutic outcome. The promise of such drugs is that they would preferentially target activated immune cells expressing eHsp90 with potentially minimal side effects, in contrast to current broadly acting RA therapies such as TNF-α inhibitors [9] [1; 14].

**Figure 6.**
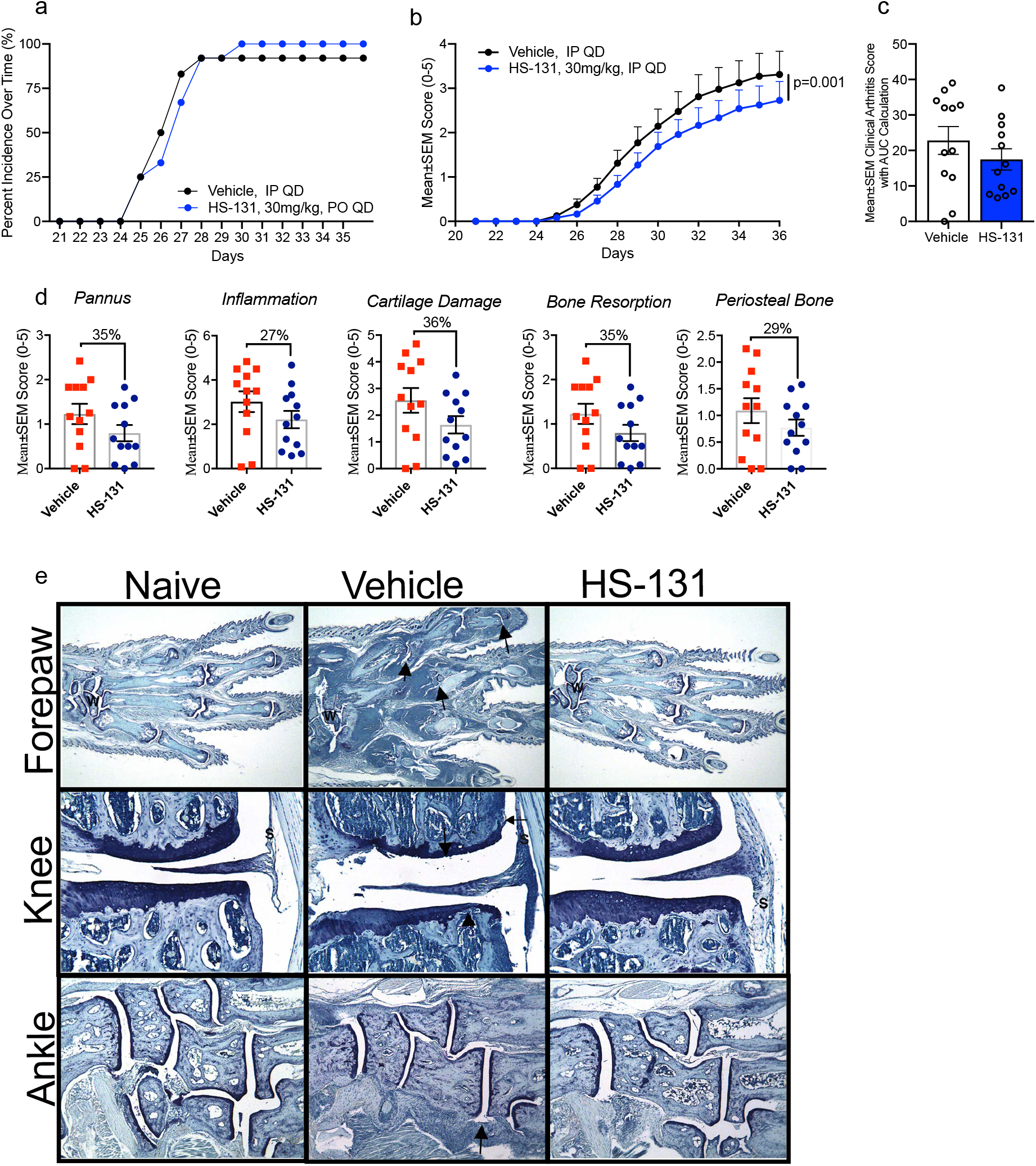
Therapeutic potential of eHsp90 targeted therapies for the treatment of auto immune diseases. (A) Animals developing disease, identified by inflammation within at least one paw (B) Mean arthritic clinical score of HS-131 30mg/kg IP, QD and vehicle IP, QD treated mice throughout the study duration N=12/group±SEM. (C) Area under the curve of HS-131 treated mice compared to vehicle N=12/group±SEM. (D) HS-131 reduced pannus, inflammation, cartilage damage, bone resorption and periosteal bone histological manifestations of CIA 36 days post disease onset N=12/group±SEM. Experiment was performed once with 12 biological replicates per group. (E) Representative photomicrographs of forepaw, knee and ankle from a disease control (Naïve), diseased vehicle treated and HS-131 treated animals. W identifies wrist, Arrows identify representative affected joints, (S) identifies inflammation.

## Discussion

Here we report that eHsp90 expression is dramatically upregulated in activated T cell populations in response to cytokine challenge or *in vivo* autoimmune activation. Thus, with respect to expression of eHsp90, activated T cell populations behave similarly to tumor cells exhibiting a malignant phenotype^10,11^. By contrast, even in a highly stressed systemic disease state, such as mouse models of RA and IBD, quiescent or fully differentiated cells do not express eHsp90 or absorb HS-131. In malignancy, the precise molecular mechanism governing expression of eHsp90 has yet to be defined. However, it has been shown that over-expression of known oncogenic protein kinases such as HER2 can induce benign tumor cells to express Hsp90 at the plasma membrane [5; 7; 20]. Our studies with IL-2 now demonstrate for the first time that eHsp90 trafficking can be actively regulated through receptor mediated signaling. This process is not merely the consequence of overexpressed Hsp90 due to a general cellular stress response. A possible role for eHsp90 in T cells is the chaperoning of CD surface proteins like CD3, CD4 and CD8, since we and others have noted loss of trafficking of these proteins to the cell surface in the presence of diffusible Hsp90 inhibitors[2]. Additionally, our *in vivo* work in animal models of autoimmune disease has shown that HS-131 is able to label sites of edema and inflammation, which potentially provides a new way to identify and monitor active inflammatory disease.

We further tested the potential of HS-131 to reduce disease activity in autoimmune diseases. Despite rapid biliary clearance, a small but statistically significant reduction and delay of symptoms was observed in the CIA mouse model of human RA. These findings suggest that longer acting eHsp90 targeted inhibitors could be effective and precise antiarthritic drugs. In this context, targeting eHsp90 may selectively remove auto-antigenic T cell populations permanently, perhaps even those involved in T cell memory. Our studies with fluor-tethered inhibitors of Hsp90, first in cancer and now immune disease models, has shown that expression of the extracellular form of Hsp90, eHsp90, is associated with rapidly proliferating cells and not fully differentiated cell populations or those preprogramed to be actively replaced such as gut epithelium. T cells are essential to our immune system’s ability to fight infections; however, many diseases confer maladaptive T cell activation including RA, lupus and IBS as well as other autoimmune diseases. If our hypothesis is correct, the unique association of eHsp90 expression with T cell activation opens up several avenues that have both diagnostic and therapeutic relevance with respect to autoimmune disease, managing transplant rejection and certain viral infections. For diagnostic purposes, with the advent of fluor-tethered Hsp90 inhibitors, one can now track T cell activation in blood samples in response to infection or in autoimmune disease. Ultimately our results suggest that further evaluation of the therapeutic potential of Hsp90 inhibitors in autoimmune disease is warranted. If one could intervene at the time of activation in certain autoimmune diseases that are associated with infection, such as type I diabetes or chronic Lyme disease, perhaps one could prevent the aberrant T cell “education” that ultimately gives rise to underlying chronic pathology associated with these infections. In situations of active ongoing chronic disease, tethered eHsp90 inhibitors carrying toxins or radionucleotides could be employed to acutely, precisely eliminate host targeting T cell populations without affecting the quiescent population. Once the inhibitor had been cleared by the hepatobiliary system, the patient would not be expected to suffer from long-term immunosuppression.

## Acknowledgements

We thank Jason Geswiki (UCSF) for providing the comprehensive list of proteins comprising the cellular chaperone machinery. This work was originally funded by grants from the NIH 5R01AI089526-05 (to TAJH) and 1RI090644-04 to J.J.K. and T.A.J.H.

## Data Availability

Data that supports the findings of this study are available from the corresponding author upon reasonable request.

## Methods

### T-cell activation

PBMC aliquots (ZenBio SER-PBMC-F) were washed in RPMI-1640 with 10% FBS and PSG and then rested overnight in a T-75 flask. The suspended cells were then transferred to a fresh flask and activated in fresh RPMI-1640/10% FBS/PSG by 3:20,000 anti-CD3 (BioLegend 317303), 3:20,000 anti-CD28 (BioLegend 302913), and 1:10,000 (30 IU/mL) human IL-2 (CST 8907SC) and incubated for up to 72 hours. In the experiments with longer activation, samples were diluted 1:3 into fresh media containing hIL-2 at 72 hours post initial dosing. Rested cells were treated similarly except they did not receive activating antibody or hIL-2.

### Activation inhibition experiment-Hsp90 antibodies

ADI-SPS-771, 9D2, and AC88 (all Enzo) were cleaned with 10kD MWCO spin filters. Cleaned antibody was added in a titration to newly activated or resting PBMCs. The cells were incubated for 72 hours and then stained for flow cytometry.

### Western Blot

Samples were prepared as described in the specific experiment and then loaded on Criterion 4-15% Tris-HCl gels (BioRad 5678083 and 5678084) and separated at 200V. The proteins were transferred from the gel to a PVDF membrane (BioRad 1620177) on ice for 1 hour at 100V. After transfer, the membranes were rinsed in 1x TBST and then blocked for 30 minutes with either 5% NFDM/TBST or 5% BSA/TBST depending on the block being used for the primary antibody staining step. After blocking, the membranes were labelled with primary by overnight incubation with the manufacturer recommended concentration and blocking agent for each antibody (Table 1). Residual primary antibody was removed by 5-minute washes with TBST (3x) and then the appropriate secondary antibody (CST 7074 and 7076) was added at 1:10,000 in 5% NFDM/TBST for 1 hour at room temperature. Upon completion of secondary stain, residual secondary antibody was removed by 5-minute washes with TBST (3x). Clarity ECL (Biorad 1705060) reagent was used to develop film.

**Table 1:**
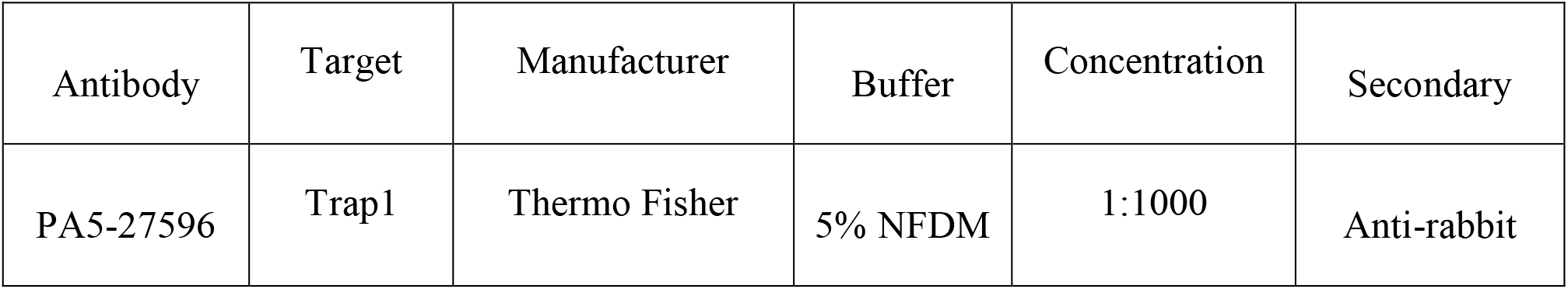
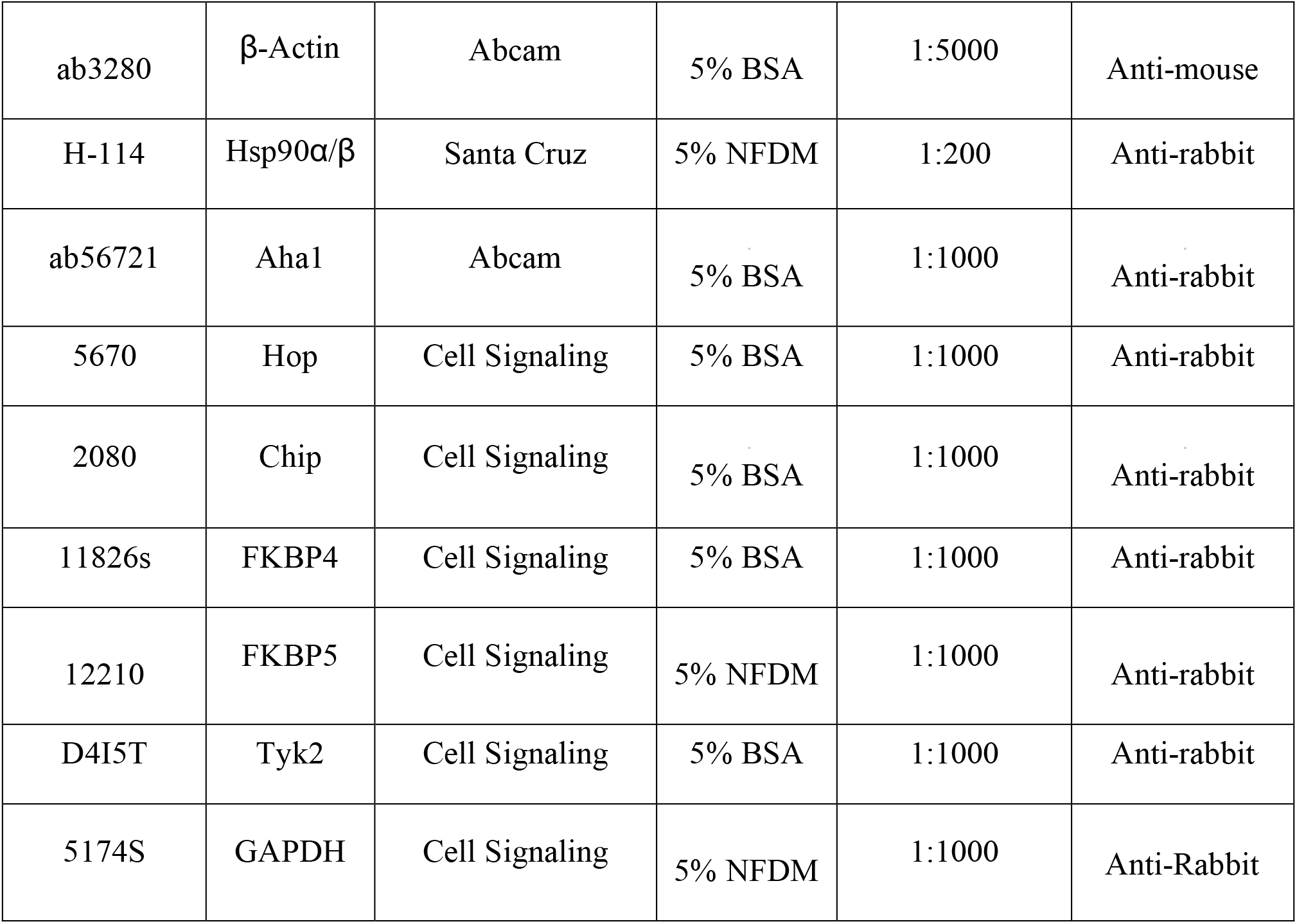
Western blot antibodies.

### Mass Spectrometry

Following silver stain of SDS-PAGE, visible bands were excised from the gel. Gel pieces were distained using 1:1 30mM potassium ferricyanide: 100mM sodium thiosulfate for 10 minutes followed by alternating washes with 25mM ammonium bicarbonate and acetonitrile. Following dehydration with acetonitrile, 25µL of porcine trypsin (Promega)at a concentration of 4µg/mL was added to the gel pieces. Following digestion, the supernatant was transferred to a second tube, and acetonitrile was added to the gel pieces to complete the extraction of digested peptides. This extract was added to the first supernatant and this combined solution, containing the extracted peptides was frozen and lyophilized. The peptides were resuspended in 5µL of 100:99:1 acetonitrile: water: trifluoroacetic acid immediately prior to spotting on the MALDI target.

For MALDI analysis, the matrix solution consisted of alpha-cyano-4-hydroxycinnamic acid (Aldrich Chemical Co. Milwaukee, WI) saturating a solution of 1:1:0.01 acetonitrile: 25mM ammonium citrate: trifuoroacetic acid. Approximately 0.15 µL of peptide solution was spotted on the MALDI target immediately followed by 0.15µL of the matrix solution. MALDI MS and MS/MS data was then acquired using the ABSCIEX TOF/TOF. 5800 Mass Spectrometer. Resultant peptide mass fingerprint and peptide sequence data was submitted to the UniProt database using the Mascot search engine to which relevance is calculated and scores are displayed.

### Flow Cytometry

The activated/rested cells were plated into a 96-well round bottom plate and pelleted at low speed. Cells were resuspended in chilled media containing titrated HS-131 or HS-198 (in 1% DMSO) and stained for 30 minutes at RT. Fluorochrome tagged inhibitors and media were washed away with a PBS wash, followed by a PBS wash containing 3% NMS. Surface antibody stains were performed in 3% NMS/PBS for 30 minutes followed by PBS washes (2x). Samples were analyzed on BD FACSCanto II flow cytometer. Permeabilized samples were prepared as previously described followed by fixation in 4% Formaldehyde/PBS overnight at 4°C. Sample analysis was performed with Flowing Software and FlowJo.

### Flow cytometry antibodies

| Antibody | Target and fluor | Manufacturer | Concentration |
| --- | --- | --- | --- |
| 300434 | CD3- BV421 | BioLegend | 1:200 |
| 302616 | CD25-AF488 | BioLegend | 1:200 |
| 423113 | Viability-BV421 | BioLegend | 1:1000 |
| 423102 | Viability-BV510 | BioLegend | 1:1000 |
| 310903 | CD69-FITC | BioLegend | 1:200 |
| 302605 | CD25-PE | BioLegend | 1:200 |
| 561105 | Ly6G-AF488 | BD Biosciences | 1:500 |
| 103139 | CD45-BV605 | BioLegend | 1:800 |
| 563415 | IA/IE-BV650 | BD Biosciences | 1:1500 |
| 103121 | CD45-AF488 | BioLegend | 1:1000 |
| 300454 | CD3-AF488 | BioLegend | 1:1000 |
| 301817 | CD14-AF488 | BioLegend | 1:1000 |
| 400625 | Rat IgG2b Isotype Control | BioLegend | 1:1000 |

### Flow on CIA mouse Cells

#### CIA Disease

Collagen was prepared at 4 mg/ml in 0.01 N acetic acid. Equal volumes of 4 mg/ml collagen and 5 mg/ml Freund’s complete adjuvant were emulsified by hand mixing with syringes for approximately 5 min, at which point a bead of this material holds its form when placed in water. On study days 0 and 21, animals were anesthetized with isoflurane and given intradermal injections of a total of 400 µg of type II collagen in Freund’s complete adjuvant at the base of the tail.

#### CIA Experimental Design

Mice were randomized into treatment groups by body weight on study day 18. Animals were treated from day 21-36 of the study with either 20 μL of HS-131 (30mg/kg) PO, QD or vehicle (DMSO) PO, QD. On study day 36, the mice were euthanized for necropsy. Clinical scores were given for each of the paws (right front, left front, right rear, left rear) on study days 21–36. 0= normal, 1= one hind or fore paw joints affected or minimal diffuse erythema and swelling, 2= two hind or fore paw joints affected or moderate diffuse erythema and swelling, 3= three hind or fore paw joints affected or moderate diffuse erythema and swelling, 4= four hind or fore paw joints affected or marked diffuse erythema and swelling, 5=entire paw affected, severe diffuse erythema and severe swelling, unable to flex digits. Experimenter was blinded from the treatment group during clinical evaluation and scoring.

#### IBD Disease

On study day 0, Balb/C mice were terminated and spleens obtained for CD4+CD45RB^high^ cell isolation. After cells had been sorted and obtained, each animal received an IP injection of at a minimum 4×10^5^ cells (200µl/mouse injections). On study day 49, a diseased mouse was treated IV with 20μl of inflammation tracer HS-131 (10nmoles). The animal was observed for 24 hours. On study Day 50, the mouse was euthanized via CO2 inhalation, frozen in liquid nitrogen using a black cryoprotectant gel (embedding medium), and sent to BioInVision for analysis.

#### CryoViz™ Imaging

Whole mice were imaged at 10.23 *μ*m × 10.23 *μ*m in-plane resolution using an Olympus MVX-10 microscope with a 1X objective and 0.63X magnification and 40 *μ*m section thickness, using the CryoViz™ (BioInVision, Inc., Cleveland, USA). CryoViz™ is a fully-automated, serial sectioning-and*-*imaging system which provides 3-dimensional, tiled, microscopic anatomical bright field and molecular fluorescence images over large fields-of-view such as a whole mouse. Images were acquired using a dual band FITC/TxRed fluorescence filter (Chroma, Inc., Rockingham, VT), a liquid crystal RGB filter and a low-noise monochrome camera. Raw images acquired by the CryoViz™ were processed to generate 3D color anatomical brightfield and molecular fluorescence volumes using the CryoViz™ Preprocessor software (BioInVision, Inc., Cleveland, Ohio). These 3D volumetric image data were then processed using the CryoViz™ 3D Visualizer software (BioInVision, Inc., Cleveland, USA) to obtain 3D reconstructed brightfield and fluorescence volume renderings and movies with 2D slice cutaway animations, showing the 2D/3D biodistribution of TxRed labeled HS131 within the mouse volume.

#### Microscopy

T cells isolated from PBMC’s were plated onto coverslips. The cells were stained with HS-131 and HS-198 at room temperature for 2 hours. Following staining, cells were fixed to coverslips with 4% formaldehyde/PBS. The fixative was then removed with PBS washes (3x) and Unbound stain was removed by PBS washes (2x) and then the coverslips were rinsed with ddH2O and affixed to slides with FluorSave (Millipore 345789). Imaging was performed on a Leica SP5 confocal microscope.

#### Immunohistological Staining

After 24-48 hours in fixative and 4–5 days in 5% formic acid for decalcification, tissues were trimmed, and processed for paraffin embedding. Paws were embedded in paraffin in the frontal plane and the knees were embedded with the patella facing down. Ankles, if left attached to the hind paw, were also embedded in the frontal plane but may be detached and sectioned in the sagittal plane for special purposes. Sections were cut and stained with toluidine blue.

#### Scores for Synovitis, Pannus Formation, Degradation of Cartilage, and Bone

##### Paw Score Criteria

0 = Normal. 0.5 = Very minimal, affects only 1 joint or minimal multifocal periarticular infiltration of inflammatory cells. 1 = Minimal infiltration of inflammatory cells in synovium and periarticular tissue of affected joints. 2 = Mild infiltration of inflammatory cells. When referring to paws, generally restricted to affected joints (1–3 affected). 3 = Moderate infiltration with moderate edema. When referring to paws, restricted to affected joints, generally 3–4 joints and the wrist or ankle. 4 = Marked infiltration affecting most areas with marked edema, 1 or 2 unaffected joints may be present. 5 = Severe diffuse infiltration with severe edema affecting all joints (to some extent) and periarticular tissues.

##### Knee Score Criteria

0 = Normal. 0.5 = Very minimal, affects only one area of the synovium or minimal multifocal periarticular infiltration of inflammatory cells. 1 = Minimal infiltration of inflammatory cells in synovium and periarticular tissue of affected synovial areas. 2 = Mild diffuse infiltration of inflammatory cells. 3 = Moderate diffuse infiltration of inflammatory cells. 4 = Marked diffuse infiltration of inflammatory cells. 5 = Severe diffuse infiltration of inflammatory cells.

##### Cartilage Damage Score Criteria

0 = Normal. 0.5 = Very minimal = Affects marginal zones only of one to several areas (knees) or joints (paws). 1 = Minimal = Generally minimal to mild loss of toluidine blue staining (proteoglycan) with no obvious chondrocyte loss or collagen disruption in affected joints/areas. 2 = Mild = Generally mild loss of toluidine blue staining (proteoglycan) with focal areas of chondrocyte loss and/or collagen disruption in some affected joints/areas. Paws may have one or two digit joints with near total to total loss of cartilage. 3 = Moderate = Generally moderate loss of toluidine blue staining (proteoglycan) with multifocal chondrocyte loss and/or collagen disruption in affected joints/areas. Paws may have three or four joints with near total or total loss. In the knee, some matrix remains on any affected surface with areas of severe matrix loss. 4 = Marked = Marked loss of toluidine blue staining (proteoglycan) with multifocal marked (depth to deep zone or tidemark) chondrocyte loss and/or collagen disruption in most joints with a few unaffected or mildly affected. In the knee, one surface with total to near total cartilage loss. 5 = Severe = Severe diffuse loss of toluidine blue staining (proteoglycan) with severe (depth to tide mark) chondrocyte loss and/or collagen disruption in most or all joints.

### Quantification and Statistical Analysis

Graphpad Prism 8 was used for statistical analysis of T cell activation, eHsp90 expression and viremia. For each analysis, total n and SEM are presented in the figure legend. An alpha of .05 was used for all statistical analysis.

