## Supplemental Figures for "Expression of extracellular Hsp90 is a molecular signature of T cell activation, providing a means to image and target T Cell activation in autoimmune disease"

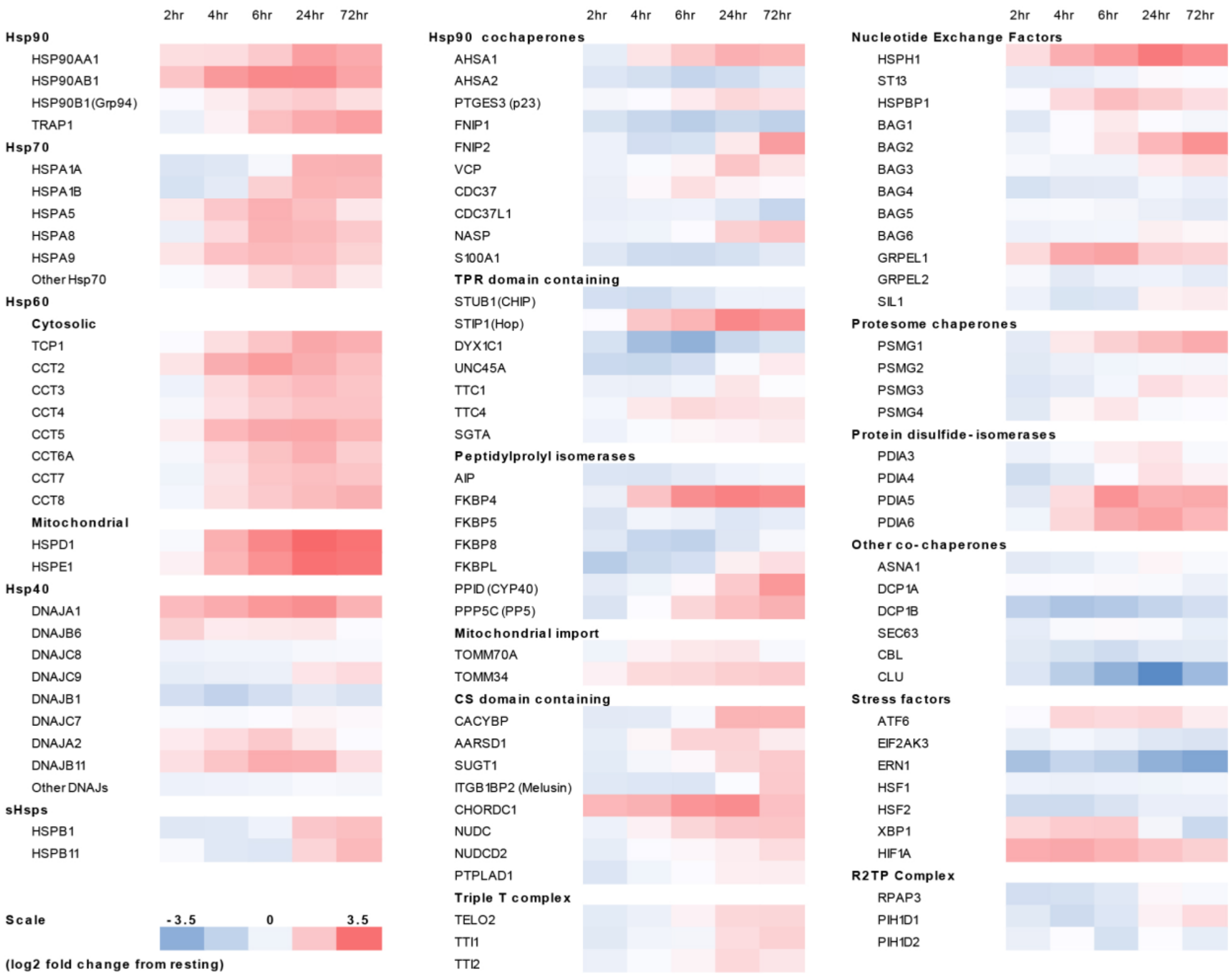


### Figure 1s.

Regulation of mRNA expression following T-cell activation.

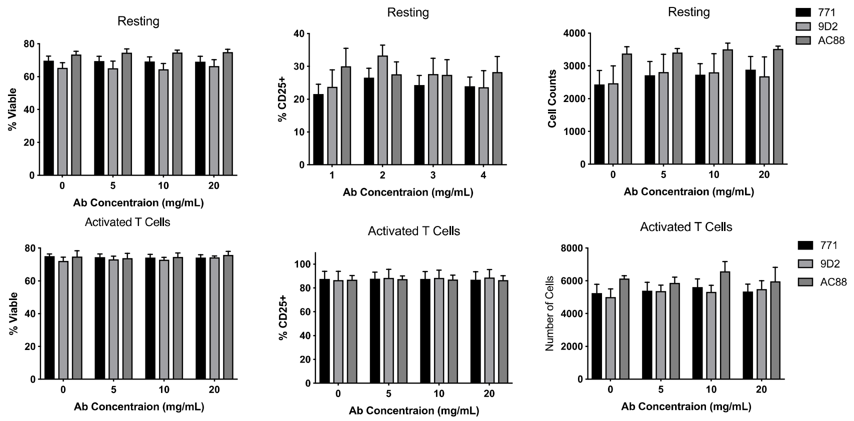


**Figure 2s.**

Effects of monoclonal Hsp90 targeted antibodies. Effects of three monoclonal Hsp90 antibodies on resting and activated T-cell viability, percentage CD25+ and number of cells.
